# A unified model of species abundance, genetic diversity, and functional diversity reveals the mechanisms structuring ecological communities

**DOI:** 10.1101/2020.01.30.927236

**Authors:** Isaac Overcast, Megan Ruffley, James Rosindell, Luke Harmon, Paulo A. V. Borges, Brent C. Emerson, Rampal S. Etienne, Rosemary Gillespie, Henrik Krehenwinkel, D. Luke Mahler, Francois Massol, Christine E. Parent, Jairo Patiño, Ben Peter, Bob Week, Catherine Wagner, Michael J. Hickerson, Andrew Rominger

## Abstract

Biodiversity accumulates hierarchically by means of ecological and evolutionary processes and feedbacks. Reconciling the relative importance of these processes is hindered by current theory, which tends to focus on a single spatial, temporal or taxonomic scale. We introduce a mechanistic model of community assembly, rooted in classic island biogeography theory, which makes temporally explicit joint predictions across three biodiversity data axes: i) species richness and abundances; ii) population genetic diversities; and iii) trait variation in a phylogenetic context. We demonstrate that each data axis captures information at different timescales, and that integrating these axes enables discriminating among previously unidentifiable community assembly models. We combine our massive eco-evolutionary synthesis simulations (MESS) with supervised machine learning to fit the parameters of the model to real data and infer processes underlying how biodiversity accumulates, using communities of tropical trees, arthropods, and gastropods as case studies that span a range of spatial scales.

## Introduction

Biodiversity is structured hierarchically across spatial, temporal, and taxonomic scales (Leibold & Chase 2019). Fluctuations of species abundances within communities operate on ecological timescales, on the scale of handfuls or tens of generations. Population genetic variation, by contrast, accumulates and degrades over timescales of tens to tens of thousands of generations (Leffler *et al.* 2012), while phylogenetic and functional diversity accumulate even more slowly, on the order of thousands to millions of generations (Uyeda *et al.* 2011). Over time, various fields have emerged to investigate processes within individual levels of organization (macroecology, comparative population genetics, macroevolution), but only recently have inroads been made to combine theory across multiple scales of space and time into a general unified model (Vellend 2010, 2016). Complicating matters, there is little consensus over whether, and to what degree, ecological interactions contribute to the structuring of ecological communities (Rabosky & Hurlbert 2015; Harmon & Harrison 2015). Likewise, the relative contributions of colonization and *in situ* speciation to the composition of community structure remains an open question (Patiño *et al.* 2017). Feedbacks across biological levels of organization are well known, yet we continue to lack a model of community assembly that accounts for such feedbacks.

Discovering universal rules that structure ecological communities is a challenging task given the difficulty of disentangling the relative influence of faster ecological mechanisms from slower evolutionary processes (Ricklefs 2004), yet a unification of theory across multiple scales will provide significant insight into the formation of biodiversity (McGill *et al.* 2019). Ecological models of community biodiversity inspired by the Equilibrium Theory of Island Biogeography (MacArthur & Wilson 1967) and the Neutral Theory of Biodiversity and Biogeography (Hubbell 2001) have focused on predicting the shape of the local species abundance distribution (SAD). As central as the SAD is to macroecology and community ecology, it is often insufficient to distinguish among different models of community assembly, particularly at equilibrium (Chave *et al.* 2002; McGill *et al.* 2007, Haegeman & Etienne 2011). Progress has been made toward linking community ecology models with population genetics (Baselga *et al.* 2013; Baselga *et al.* 2015; Vellend 2005), however, current theory either lacks an explicitly population genetic foundation (Vellend 2005), or considers genetic variation only of a focal species (e.g. Laroche *et al.* 2015). A great deal of work has been done to incorporate phylogenetic information with abundance data to make inferences about community assembly processes (Webb *et al.* 2002, Jabot & Chave 2009). While such approaches make useful predictions, they are predicated on assumptions of equilibrium within the local community, and also assume that the phylogeny is a reliable proxy for functional trait diversity (Cavender-Bares *et al.* 2009, Mayfield & Levine 2010). There have been other efforts to unify different time-scales with mechanistic eco-evolutionary models of assembly. For example Cabral *et al.* (2019) unify population-level and evolutionary timescales to investigate the dynamic relationship between community age, competition, and local richness. Likewise, Pontarp *et al.* (2019a) develop a trait-based, spatially explicit eco-evolutionary model to make inferences about prey and predator niche width with potentially diverse data types. Incorporating temporal dynamics can help to distinguish among processes (Azaele *et al.* 2006; Chisholm & O’Dwyer 2014; Jabot *et al.* 2018; Kalyuzhny *et al.* 2015; Nee 2005; Ricklefs 2006), yet current theory fails to generalize across levels of biological organization. Adding more axes of data to process-based models without increasing model complexity at the same rate is therefore a necessary advance to break this many-to-one mapping of hypotheses to observation (McGill *et al.* 2007; Leibold & Chase 2017).

The massive multi-dimensional datasets that continue to emerge from high-throughput biodiversity investigations applying community-wide surveying techniques such as eDNA (Deiner *et al.* 2017), metabarcoding (Andújar *et al.* 2018; Dopheide *et al.* 2019), and remote-sensing technologies that can directly infer trait data (Cavender-Bares *et al.* 2017), are therefore timely. However, the challenges associated with moving beyond descriptive approaches of interpretation and inference have limited broader understanding of processes generating biodiversity patterns (but see Bohan *et al.* 2017; Derocles *et al.* 2018). Historically there have been two general approaches to investigate the evolutionary and assembly processes underlying the patterns we observe in nature: 1) “process-first” approaches that use first principles to derive generative mechanisms to make predictions of a single data type under the assumptions of an idealized community (Gavrilets & Vose 2005; Rosindell *et al.* 2012; Marquet *et al.* 2014); and 2) “pattern-first” approaches that reveal aggregate differences in macroecological patterns from real world systems across a range of spatial and temporal scales (Craven *et al.* 2019; Keil & Chase 2019; Ricklefs & Bermingham 2001; Rominger *et al.* 2016; Wagner *et al.* 2014). Recent advances in simulation-based inference under increasingly complex models provide a third option of unifying multiple processes and multiple data types across different scales (e.g. Overcast *et al.* 2019; Pontarp *et al.* 2019b). This unified model of community assembly, which accounts for fundamental processes underlying biodiversity across spatial and temporal scales, could be used to make predictions about multiple axes of biodiversity data that include species richness and abundances, distributions of species genetic diversities, and trait variation. Several studies have recently shown that such complex biological models and resultant high-dimensional data can be tractable within a machine learning framework (Schrider & Kern 2018; Sheehan & Song 2016), providing a robust inference procedure for simulation-based interrogation of empirical data.

Here we introduce the Massive Eco-evolutionary Synthesis Simulations (MESS) model, building upon classic community ecology theory (Hubbell 2001; Leibold & Chase 2017; MacArthur & Wilson 1967; Vellend 2016) to produce a new mechanistic eco-evolutionary model of community assembly for making dynamic joint predictions of observed data. MESS integrates ecological models of community biodiversity, comparative population genetics, and community phylogenetics, with an explicit focus on incorporating microevolution and ecological interaction processes, which are often underrepresented in mechanistic models (Leidinger & Cabral 2017). MESS can simulate community assembly under an array of models across a continuum of evolutionary scenarios (niche versus neutral and evolved versus assembled). These simulations generate community-scale distributions of abundance, genetic variation, and trait values which are summarized using a novel combination of statistics that capture the variation within and among these biodiversity data axes. We combine summary statistics from numerous simulations with supervised machine learning methods to test an array of competing models and to estimate model parameters relevant to understand complex histories of community assembly and evolution. We perform extensive simulation-based cross-validation analyses to explore precision and accuracy of model inference. Finally, we apply the model to four empirical datasets representing different taxonomic and spatial scales: two arthropod communities with varying dispersal capacity from Mascarene islands of different ages (Emerson *et al.* 2017; Kitson *et al.* 2018); plot level sampling of Australian tropical forest trees (Rossetto *et al.* 2015); and archipelago-scale sampling of micro-endemic terrestrial gastropods from the Galapagos Islands (Kraemer *et al.* 2019; Triantis *et al.* 2016). Most empirical communities obtain approximately equilibrium structure, with varying degrees of inferred neutrality, however, the tree community was far from equilibrium, possibly because of higher rates of local turnover and stronger species-specific environmental filtering.

## Methods

### Metacommunity composition

The MESS model comprises three components summarised in Figure 1. The metacommunity is modelled as a regional pool which is very large and fixed with respect to the timescale of the assembly process in the local community. It consists of a global phylogeny relating all species, along with species abundances, and trait values evolved along the phylogeny. The global phylogeny is produced by simulating a constant birth-death process with fixed speciation (λ) and extinction (λ · ∊) parameters, until the desired number of species (*S_M_*) is reached (*TreeSim* v2.4; Stadler 2019). Next, we simulate a Brownian motion model of trait evolution on the phylogeny with a root value of 0 and a rate of σ^2^ (*ape* v5.3; Paradis *et al.* 2004). Traits evolve following a Brownian motion process in the metacommunity, rather than an Ornstein–Uhlenbeck process (Butler & King 2004), because we assume species in the metacommunity are not exposed to constraints imposed by the local environmental conditions. Additionally, we do not model intraspecific trait variation, on the assumption that trait values represent the mean phenotype of each species. Finally, the species abundances are sampled from a log-series distribution parameterized by the total number of species (*S_M_*) and the total metacommunity size (*J_M_*).

**Figure 1:**
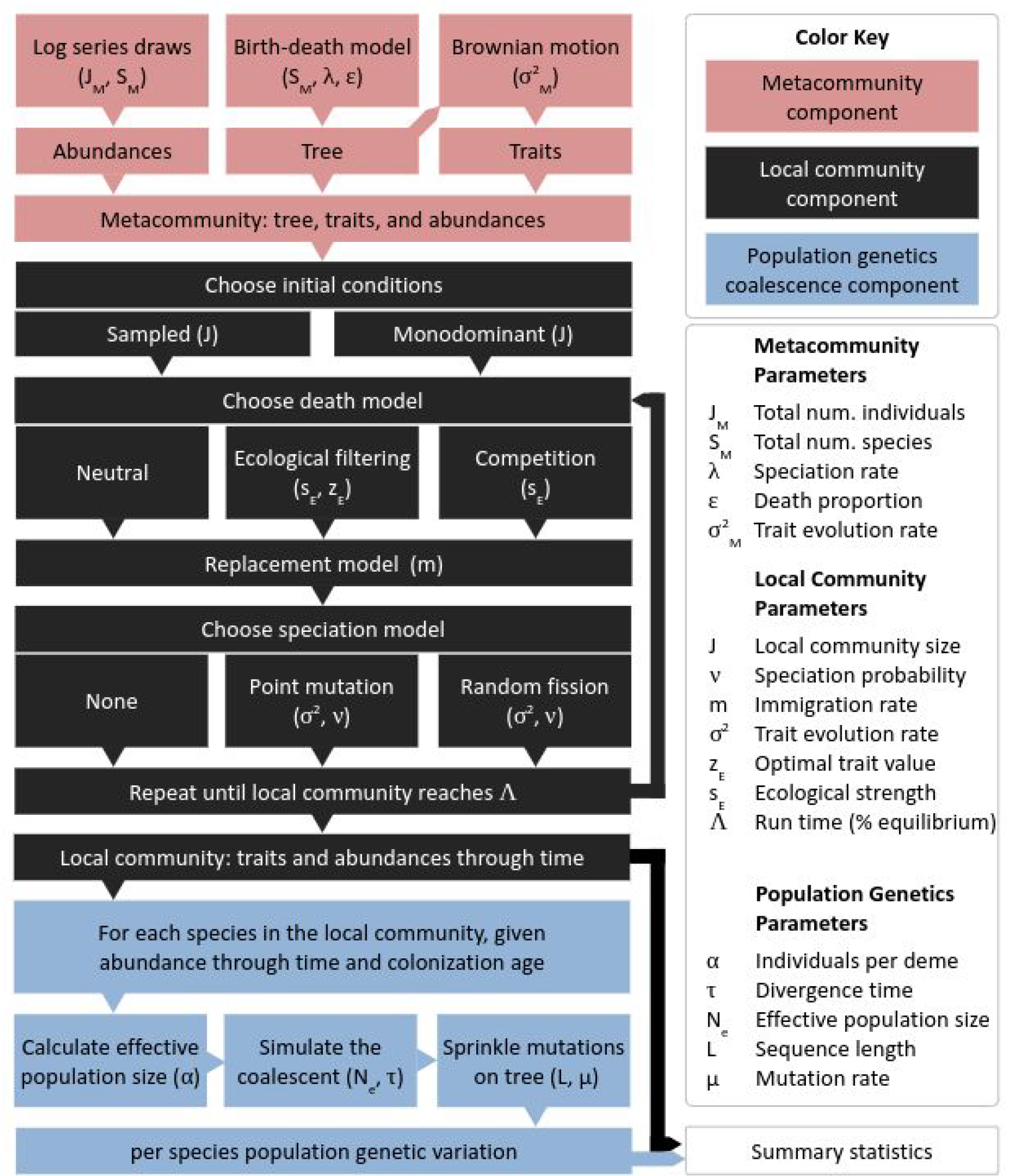
Conceptual diagram illustrating the three primary components of MESS. The metacommunity component (red) encompasses of a global phylogeny relating all species, along with species abundances and trait values evolved along the phylogeny. The local community component (black) involves a forward-time process during which a local community assembles by birth, death, immigration, and local speciation. The population genetic component (blue) generates backward-time coalescent simulations per species which are parameterized contingent on the abundance history and colonization time generated by the forward-time component to approximate the accumulation of genetic diversity. Each box illustrates a sub-component of the model, and indicates the parameter(s) which determine the behavior of each sub-component. Arrows between sub-components indicate information flow through the process.

### Local community dynamics

The foundations of the community dynamics underlying MESS are based on the joint neutral model of abundance and genetic diversity described in Overcast *et al.* (2019). Briefly, we simulate an individual-based model of community assembly inspired by the ecological neutral theory of Hubbell (2001), with assembly in a local community proceeding by a process of birth, death, and colonization from the metacommunity (following Rosindell & Harmon 2013). Departing from the previous model, MESS local community dynamics can range from fully neutral (species traits have no effect), to various degrees of non-neutrality determined by the magnitude that species traits influence individual death probability (δ) through competition or environmental filtering. Following Ruffley *et al.* (2019), we based our environmental filtering and competition models on a functional relationship common in coevolutionary models which relates trait-based interactions with the probability of persistence in a community, scaled by the ecological strength (*s_E_*; Lande 1976; Nuismer & Harmon 2015; Andreazzi *et al.* 2017). The *s_E_* parameter determines either the strength of species-species competitive interactions or species-environment filtering interactions depending on whether a competition or filtering model is specified. Calculated death rates per species are normalised to provide a vector of death probabilities that weight the random sampling of which individual will die in each time step according to a multinomial distribution (see Supporting Methods).

As a first approximation, we implement a point mutation speciation process (Hubbell 2001), although other modes could be incorporated in future versions of the model (Rosindell *et al.* 2010; Haegeman & Etienne 2017). Speciation is implemented phenomenologically and takes place with probability ν upon each birth event. Upon each speciation event, the new individual is assigned a unique species identity, and its prior species identity is recorded as the parental species for purposes of building the local phylogeny. The descendant species receives a trait value sampled from a normal distribution centered on the parent species’ trait value and with variance equal to σ^2^_M_ /(λ + λ · ε), which is the expected variance of trait differences between parent and offspring species in the metacommunity.

### Population genetics component

Following Overcast *et al.* (2019), the forward-time histories of colonization and abundance changes through time per species are used to parameterize backward-time coalescent models with immigration for each species (Kelleher *et al.* 2016) to generate sampled local nucleotide diversities (π; Nei & Li, 1979). For reasons of computational efficiency, and to achieve a realistic scale in terms of numbers of individual organisms, we use a scaling parameter (α) to specify the number of individuals per deme, thus the total number of organisms in the local community is given by *J* · α. This notion of demes, or ‘cohorts’, groups of individuals that perform the same actions at the same time, is conceptually similar to that of Harfoot et al. (2014). We use the forward-time frequency of colonization events (scaled to number of colonizations per generation) for each species to parameterize the migration probability in the coalescent of colonization/divergence with ongoing immigration. Given an observed dataset, coalescent simulations match the observed sample sizes of each species for which DNA sequence data was obtained with regards to numbers of individuals per species and length of sequence.

### Summary statistics

We specify an hierarchical structure of summary statistics for each target data axis: species abundances, population genetic variation, and trait values. First, several relevant summary statistics are calculated per species, for each of the data axes. Next, each species-level statistic is aggregated and community-scale summary statistics are calculated per axis of data, capturing information about the distribution of the statistic across the community. We include as summaries the first four moments of each community-wide distribution, as well as pairwise Spearman rank correlations among all data axes. For correlations involving the trait axis, we consider the absolute value of the difference between the species trait and the local trait mean as the trait variable. We also calculate the differences between regional and local values of trait mean and standard deviation (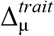 and 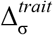 respectively). Additionally, we utilize a framework of generalized Hill numbers as community-scale summary statistics, to quantify the shape of each distribution (Chao *et al.* 2014; Gaggiotti *et al.* 2018). In order to distinguish between these diversity metrics when calculated on distributions of different data axes we will refer to the Hill number of order *q* for abundance data as *^q^D*, for genetic data as *^q^GD*, and for trait (functional) data as *^q^FD*. For simplicity, throughout the manuscript we will refer to Hill numbers calculated on distributions of each data axis as abundance, π, and trait Hill numbers.

### Model behavior

We simulated communities under a range of parameter values to understand how different model processes affect the distributions of community-scale data, and whether the summary statistics capture information to discriminate among various alternative models. Given that the MESS model is dynamic in time, we controlled for this by running each simulation to the same fixed point in the assembly process. We quantified this point as the proportional approach to equilibrium (Λ) and fixed this parameter at 0.75. This value is measured as the fraction of information about the initial state of the local community which is no longer present in the current state (see Overcast *et al.* 2019 for a full treatment of this parameter). We allowed ν to take one of three values corresponding to no-, low- and high-speciation (0, 5.10^−4^, and 5.10^−3^ respectively). We generated 10,000 simulations for each assembly model (neutral/filtering/competition) using fixed parameter values of intermediate magnitude (see Table S2 for simulation parameters). We also investigated how summary statistics of different assembly model types vary through time. To this end, we generated 10,000 simulations for each assembly model, sampling communities at different stages of the assembly process (Λ ~ U[0,1]; see Table S3 for simulation parameters). These simulations used fixed parameters of intermediate magnitude, allowing only ν to vary taking one of three values as above.

### Machine learning inference and cross-validation

The MESS package includes an automated multi-stage machine learning (ML) inference procedure, complete details of which are available in the supporting materials. Briefly, the MESS ML classification and regression procedures can be performed with a number of ensemble learning strategies including random forest (Breiman 2001) and gradient boosting (Friedman 2001). We quantify model uncertainty on parameter estimates as prediction intervals (PIs) using a quantile regression approach (Meinshausen 2006), and we implement posterior predictive simulations to assess the goodness of fit of the model to the observed data (Gelman 2003). Unless otherwise indicated, all ML algorithms are implemented in python using the architecture of *scikit-learn* (v0.20.3, Pedregosa *et al.* 2011).

We explored the power, accuracy, and bias of the ML inference procedure to classify community assembly models and estimate parameters using simulation experiments and cross-validation (CV). To evaluate assembly model classification, we generated 10,000 simulations per model class (i.e. neutral/filtering/competition) and fixed all MESS parameters at intermediate values, varying only the size of the local community (*J*) and the local speciation probability (*v*) (see Table S4 for simulation parameters). To quantify the accuracy and bias of MESS parameter estimation utilizing an ML ensemble method regression framework, we generated 10,000 community simulations per assembly model class while varying several parameters of interest (α, *J*, *s_E_*, *m*, *v*, and Λ) using log-uniform or uniform prior distributions (see Table S5 for parameters). ML estimator performance was then investigated using a K-fold CV procedure whereby simulations were split into training and testing sets, with the model being iteratively trained on each K-fold and performance being evaluated as minimized CV prediction error on the held out training set. Classifier model adequacy was quantified by the percent error rate of misclassification, and regression model accuracy was quantified by the explained variance and R^2^ (coefficient of determination) regression scores.

### Empirical examples

As case studies, we selected four systems that occupy different spatial scales and likely occupy different locations on the continua of dispersal, speciation, ecological drift and non-neutrality. Each system has some combination of community-scale data available for two of the three axes which can be considered by the model. In this way we hope to demonstrate the power of MESS across taxonomic and spatial scales, using data availability scenarios that might be encountered by empirical biologists in the present or very near future. These systems are: 1) spiders from Réunion island with abundances collected from ten 50 m × 50 m plots and 1282 individuals sequenced for one ~500bp mtDNA region (COI; Emerson *et al.* 2017); 2) weevils from two Mascarene islands (Réunion and Mauritius) which have been densely sampled for abundance and sequenced for one mtDNA region (~600bp COI) at the community-scale (Kitson *et al.* 2018); 3) three subtropical rain forest tree communities scored for multiple continuous traits and shotgun sequenced for whole cpDNA (Rossetto *et al.* 2015) and; 4) Galapagos snail communities collected from all major islands, sampled for one mtDNA region (~500bp COI; *Kraemer et al.* 2019) and scored for two continuous traits (Triantis *et al.* 2016). For each empirical dataset we conducted 10,000 simulations of each assembly model class and generated abundances, trait values, and genetic variation corresponding to genomic regions with identical numbers of base pairs under an infinite-sites model at a rate sufficient to generate diversity similar to the empirical data (see Supporting Methods for precise empirical data curation and simulation procedures). We then conducted a round of ML model selection, parameter estimation, and quantile regression to generate parameter estimates and PIs. Finally, we implemented posterior predictive simulations to assess goodness of fit of the selected model and parameters to each of the observed datasets.

## Results

### Model behavior

Simulations generated under different community assembly models produced markedly different distributions of community-scale data and summary statistics. First we considered one static point in time (at Λ = 0.75; Fig. 2). Neutral simulations generated communities with higher species richness, more even distributions of abundance as summarized by the normalized *^q^D* values, and higher mean and standard deviation of π values. Filtering and competition models were largely indistinguishable in terms of abundance and genetic diversity, with distributions of species richness, and mean and standard deviation of the population genetic statistics broadly overlapping (Fig. 2). Distributions of statistics related to trait values showed more nuanced and variable behavior, obtaining characteristics that differ between the three models. There was little distinction between models in terms of distributions of difference in local and metacommunity mean trait values 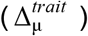, with the exception that filtering models produced more variable results. However, distributions of local and metacommunity difference in trait standard deviation 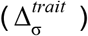 varied considerably among models, with competition tending to yield negative values (more variation locally than in the metacommunity), filtering producing positive values (less variation locally in the metacommunity), and neutral models producing values centered on zero. This pattern is borne out in Fig. 2, which illustrates the standard deviations of trait values increasing with competition, and decreasing with filtering, with respect to neutral models. The trait diversity values (*^q^FD*) tended to be slightly higher for neutral models.

**Figure 2:**
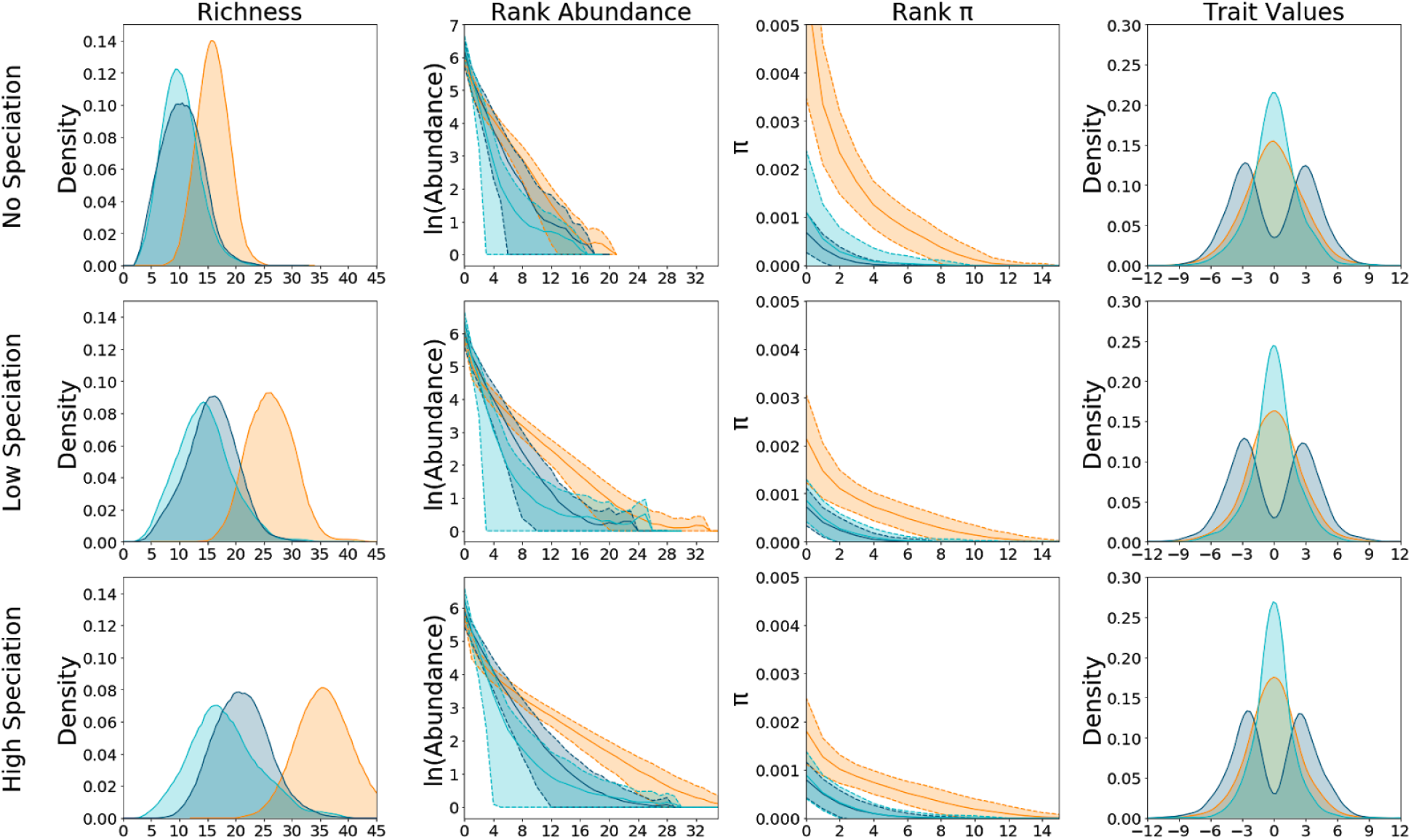
Effect of varying speciation rate and community assembly model on summary statistics. Species richness, rank abundance, rank genetic diversity, and rank distributions for 1000 simulations generated under neutral (orange), competition (dark blue) and filtering (aqua) scenarios with time fixed at 500 generations. From bottom to top, rows of panels correspond to simulations with high (ν = 0.005), low (ν = 0.0005) and no (ν= 0) speciation. In the left column of panels kernel density plots indicate the distribution of richness across simulations. In the rank plots (center two columns of panels), thick lines indicate average rank values and shaded areas show plus and minus one standard deviation. The right column of panels shows kernel density plots of zero-centered trait distributions.

Next, we investigated the temporal dynamics of MESS community histories (Fig. 3). Again, species richness in neutral models tended to exceed that of the non-neutral models throughout the entire community assembly process. In general, a low rate of local speciation produced a slight increase in richness and Hill numbers for neutral simulations, whereas a high rate produced dramatic increases in these metrics for all simulation scenarios. Between non-neutral models, richness and Hill numbers for competition were, on average, always greater than those of filtering models across all time points, with differences increasing with *v*. For neutral models, *^q^D* tended to slowly increase monotonically through time, whereas *^q^GD* initially increased quickly with community-scale genetic diversity accumulating more slowly in later stages of assembly. Increasing *v* increased the average maximum *^q^GD* for non-neutral models, but in these simulations this maximum value tended to saturate very early, with little change through time. *^q^FD* demonstrated a more dynamic temporal trajectory. Broadly, the relationships among the trait Hill numbers mirrored those of the abundance and π Hill numbers, with neutral models obtaining the highest, filtering the lowest, and competition somewhat intermediate values, and a trend of increasing values through time. However, one key difference in *^q^FD* is that early-stage communities display relatively high values, with values decreasing as Λ increases from 0 to ~0.2, and then showing an increasing trend as Λ proceeds from 0.2 to 1.

**Figure 3:**
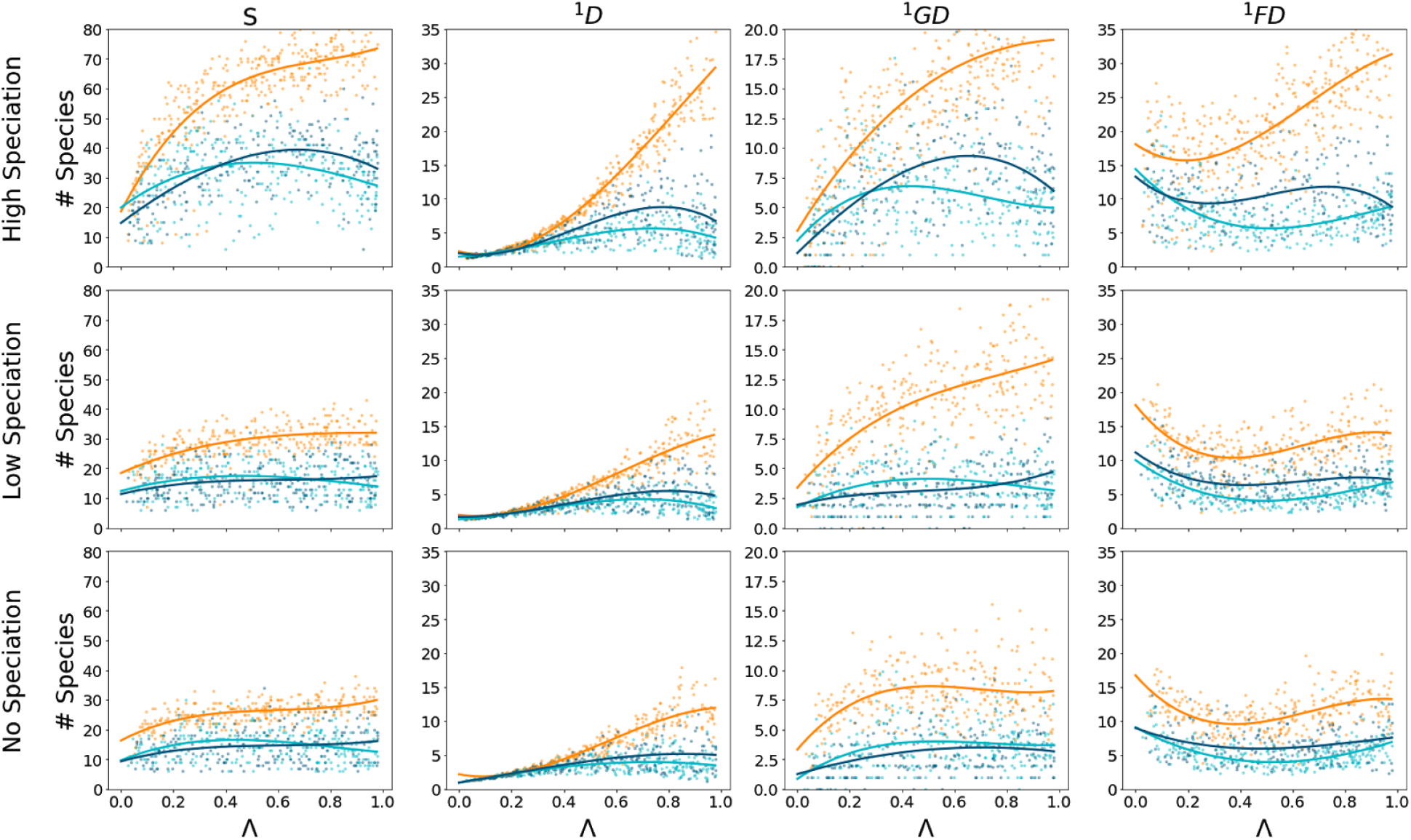
Community summary statistics through time for neutral and non-neutral models. This plot depicts the temporal change in select summary statistics for the three focal community assembly models at three different speciation rates: No, Low, and High corresponding to ν = 0, 0.0005, 0.005, respectively. Community assembly models depicted are neutral (orange), filtering (aqua), and competition (dark blue). Each subpanel shows the resultant summary statistic for 1000 simulations equally spaced through time for each model class. Simulated values are depicted as points, and a least squares polynomial is fit to better illustrate the trajectory. The far left column of panels illustrate species richness on the y-axes (S). The y-axes of the remaining columns illustrate the Hill number of order 1 for abundance, genetic diversity, and trait values, respectively.

### Model selection ML cross-validation

ML model classification prediction error reached a minimum value with *J* of 1000 for all model classes and all evaluated feature sets (Fig. 4; mean error rate 0.16). Prediction error was slightly higher for small *J* (mean error rate 0.19), and did not improve dramatically when increasing *J* from 1000 to 2000 (mean change in error rate −0.02). Neutral simulations were more accurately classified than non-neutral simulations across all feature sets and *v* values (mean error rate 0.05 and 0.18 respectively). ML classifiers trained using summary statistics from all data axes were most accurate; however, including trait information along with just one other data axis (either π or abundance) produced classification error rates close to models trained on the full suite of summary statistics. ML classifiers trained using only summary statistics related to abundance and π produced accurate classification of neutral simulations (mean error rate 0.05), but failed to distinguish between the two non-neutral models (error rate > 0.4). Importantly, in this condition the predicted model class for non-neutral simulations was overwhelmingly the alternative non-neutral model and rarely the neutral model. For example, simulations under a competition model were misclassified as filtering (0.35) with a much higher rate than neutral (0.08).

**Figure 4:**
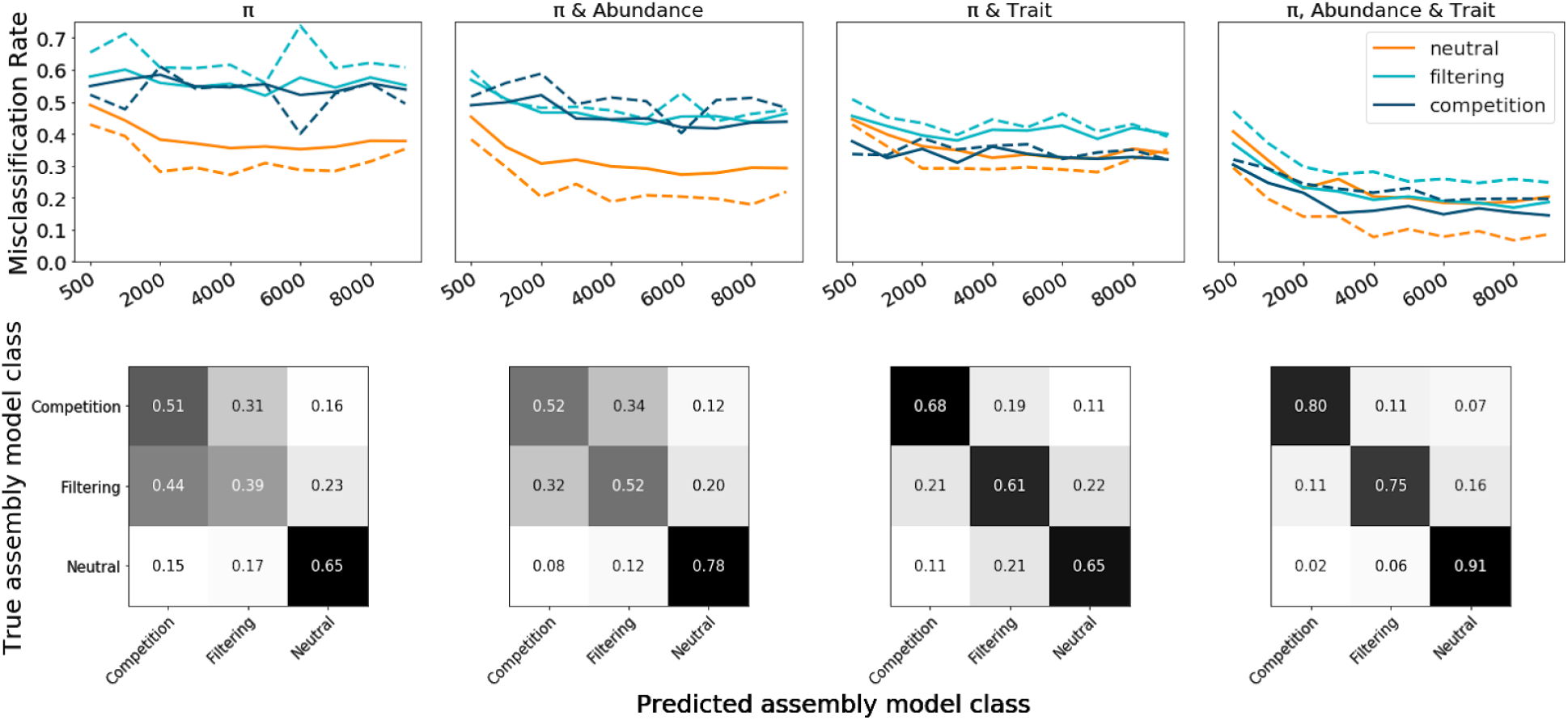
Machine learning classification error rates and confusion matrices. The top row shows random-forest misclassification error rates given different combinations of available data axes for varying sizes of local communities (J). Data axes used for each suite of simulations are indicated along the top of the figure. The x-axis indicates increasing sizes of J, from 500-10,000 in regular intervals. The y-axis indicates probability of assembly model misclassification, averaged over 1000 simulations per model class for each J (i.e. lower values indicate more accurate classification). In the figure, orange shows neutral simulations, aqua shows filtering, and dark blue shows competition. Solid lines indicate precision and dashed lines indicate recall. The bottom row shows confusion matrices depicting detailed model misclassification rates for data availability scenarios given J values between 9000 and 10,000. In these figures, values on the diagonals indicate the proportion of accurately classified simulations for each model class. Off-diagonal values indicate misclassified simulations.

### Parameter estimation ML cross-validation

Cross-validation explained variance and R^2^ regression scores for model parameter (α, *J*, *s*_*E*_, *s*_*C*_, *m*, *v*, and Λ) estimation were broadly congruent and positive in almost all cases, indicating that the simulated and estimated parameter values were correlated (in some cases highly so). For neutral simulations Λ had the highest R^2^ (0.963) and *s_E_* the lowest (−0.037), with most parameters having moderate R^2^ values (e.g. α = 0.567; *m =* 0.685; Fig. 5). The small R^2^ for *s_E_* is expected given that neutral simulations should have no information about strength of environmental interactions. Estimates of small to moderate values of *m* and *v* were accurate, but larger values tended to be underestimated. ML parameter estimation for simulations of filtering and competition models obtained improved accuracy to estimate *s_E_* (R^2^ = 0.146 and R^2^ = 0.287, respectively); however, R^2^ values for other parameters were reduced with respect to the neutral simulations (Figs. S1 & S2). Both non-neutral models produced diffuse estimates of α (R^2^ = 0.205 and R^2^ = 0.258) and *J* (R^2^ = 0.398 and R^2^ = 0.448). The most significant difference between the non-neutral models concerned estimates of Λ. Under competition scenarios, Λ estimates were precise but upwardly biased between Λ = 0 and 0.5, with increasing variance between Λ = 0.75 and 1. Under filtering scenarios, Λ estimates were only accurate for values close to Λ = 0.5, with decreasing accuracy as Λ moved away from this value in either direction.

**Figure 5:**
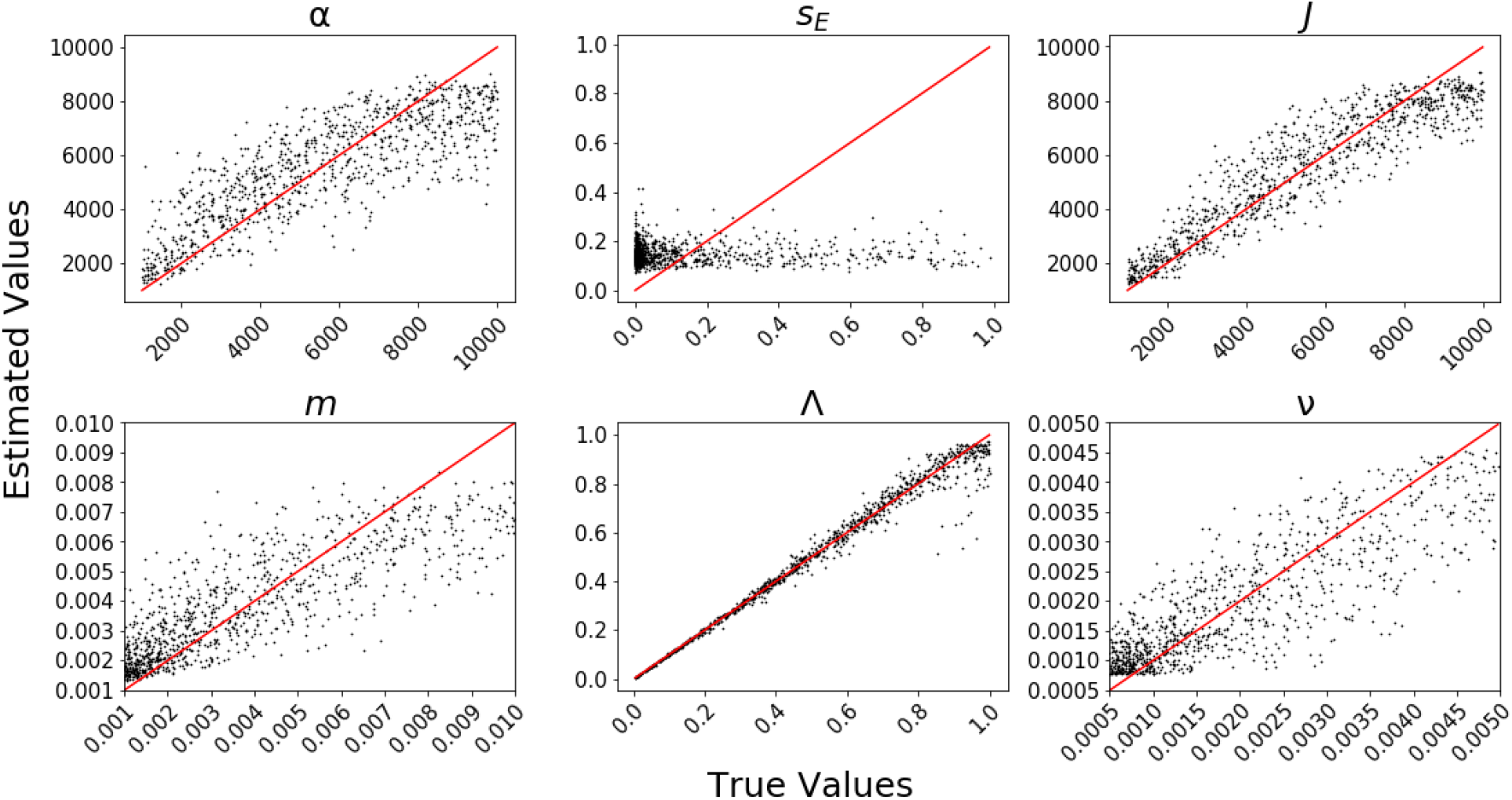
Machine learning cross-validation parameter estimation. 1000 parameter estimation cross-validation (CV) replicates using neutral community assembly model simulations and summary statistics from all data axes. True parameter values are on the x-axes and the corresponding point estimates are on the y-axes. A parameter that is well estimated will have CV results that fall on or around the identity line (depicted in red). Note that ecological strength (*s_E_*) has no impact on neutral simulations, which produces the poor CV performance in estimating this parameter.

### Empirical examples

The ML classification procedure identified the neutral model as the most probable for all three Mascarene arthropod communities (Fig. 6a), with considerable support for neutrality of the Reunion spider community (predicted class probability 0.939), and more equivocal class probabilities for Mauritius and Réunion weevil communities (0.566 and 0.53, respectively). The most important features for classification were *^1^D*, standard deviation and mean of π, *^2^D*, and *^4^D* (accounting for 44% of relative importance of all retained features).

**Figure 6:**
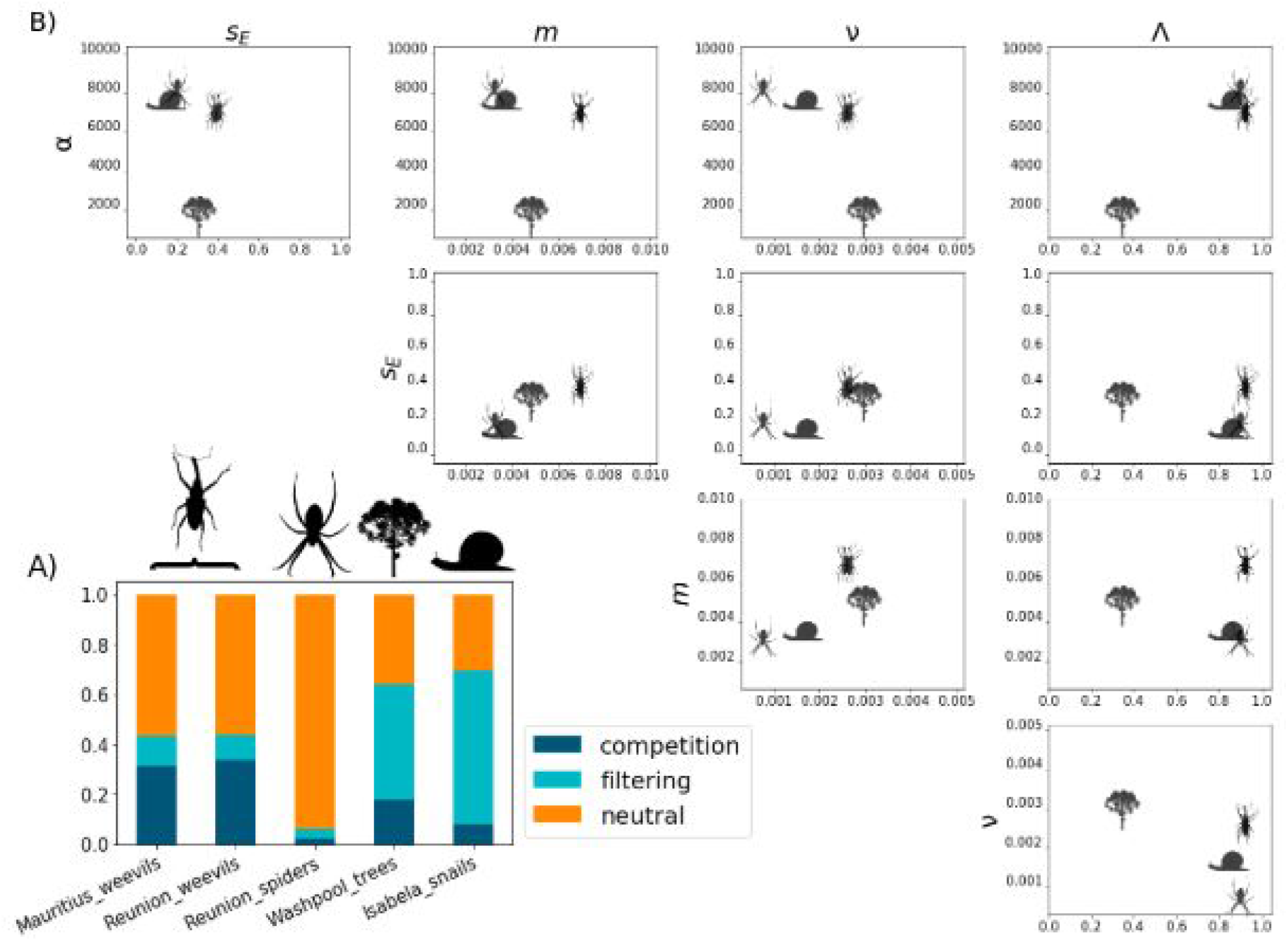
MESS empirical analysis. Empirical classification and parameter estimation of five local communities including snails, tropical trees, and island arthropods. Panel A) depicts machine learning classification probabilities for each empirical community for three focal community assembly models. The proportion of color within each bar represents the proportional predicted model class for neutrality (orange), environmental filtering (aqua), and competition (dark blue). Panel B) depicts pairwise estimates of five different model parameters under the best classified model for each local community dataset. The value along each parameter axis is indicated by the position of the representative icon. Parameters depicted include number of individuals per deme (α), ecological strength (*s_E_*), migration rate (*m*), local speciation probability (ν), and fraction of equilibrium (Λ).

The ML classification procedure identified environmental filtering as the most probable model for all tree and snail communities, with highest support for the snails (mean predicted class probability 0.698), and weaker support for the trees (mean probability 0.440). Combining filtering and competition predicted class probabilities indicated the average probability of non-neutrality for the trees was 0.633, and for the snails was 0.865. Feature importance values for classification using axes of trait and genetic data were broadly diffuse across the retained summary statistics, with 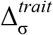 accounting for 11% of relative importance of all retained features, and the remainder accounting for 5% or less.

The ML regression procedure for parameter estimation indicated that the selected empirical datasets occupied a broad swath of parameter space (Fig. 6b; Table S6). Empirical PIs were quite varied, with some parameter estimate PIs spanning the width of the prior, while the PI of other parameters were narrow, a result which is consistent with CV results. The tree communities had small α estimates with narrow PIs (mean α = 1423; 1019-2481 95% PI), when compared to the arthropod and snail communities, which had larger α estimates (e.g. Mauritius weevil α = 7107; 3497-9831 95% PI). ML estimates of Λ were more varied, with the weevil and spider communities approaching or reaching Λ = 1, snail communities having more intermediate Λ, and tree communities having the lowest values (< 0.4 in all cases). Estimates of *m* and ν displayed an idiosyncratic pattern, with spider and snail communities having low estimated values for both, weevils having high estimated values for both, and trees having high ν and low *m* estimates. Consistent with the CV experiments, ecological strength (*s_E_*) was the most difficult parameter to estimate, in the sense that all estimates were close to the mean of the prior, and PIs spanned the majority of the prior range. Posterior predictive simulations indicated a good fit of the estimated parameters to all empirical datasets (Fig. S3).

## Discussion

We have described an individual-based mechanistic model of community assembly, the MESS model, which unifies the key processes underlying the dynamics of local accumulation of biodiversity across multiple timescales: dispersal, stochastic drift, deterministic competition/filtering, and speciation (Vellend 2010, 2016). The MESS model integrates these processes in an hierarchical framework to make multi-dimensional predictions using summary statistics that capture information both within and among the various axes of data. Simulation experiments show that neutral models have elevated *S*, *^q^D*, *^q^FD*, and *^q^GD* compared to filtering and competition models across all except the earliest time points (Fig. 3). This is a direct result of the ecological equivalence of individuals in neutral models generating communities with lower species dominance. In a similar fashion, for non-neutral models, species that are more fit survive preferentially and increase in abundance, reducing evenness in the community and causing *^1^D* to plateau at a low level, though it should be noted MESS does not implement negative density dependence and this is an avenue for future research. Increased speciation rate has little impact on *^1^D* in the neutral case because ecological equivalence confers no cost or benefit to offspring species, whereas in non-neutral models new species inherit ancestral trait values with small perturbation. In these conditions increasing speciation rate increasingly favors the evolution and accumulation of small clades of species that have ecological advantage, causing a concurrent reduction in *^1^D*.

Overall, we find that any two of the three data axes are sufficient to accurately identify the relative strength of neutral versus non-neutral processes in local community assembly, and that including trait information allows discrimination between which of the non-neutral processes are more important in driving the local patterns of biodiversity (Fig. 4). These results should be robust to values of *s_E_* that generate moderate to strong non-neutrality (i.e. s_E_ ≥ 1), with a corresponding increase in misclassification rate as *s_E_* approaches 0. More generally, using any two data axes always resulted in improved classification accuracy when compared to using a single axis alone. Furthermore, our results highlight the flexibility of MESS to mask unobserved summary statistics such that inference can be made from a wide variety of high-throughput biodiversity surveys across different spatial scales and data availabilities.

The empirical communities we chose to evaluate represent both a variety of available data axes, and a range of perceived dispersal limitation, with Galapagos snails being the most dispersal-limited, the Australian trees being least limited, and the Mascarene spiders and weevils somewhat intermediate. The results from the Reunion spider community (classified as neutral with Λ approaching 1, *m* high and *v* low) are consistent with a late-stage community that is structured primarily by colonization and ecological drift (Barabás *et al.* 2013; Vergnon *et al.* 2012). Both weevil communities had similarly high estimates of Λ, but higher estimated *v*, and less clear support for classification as neutrally evolving. The snail communities were classified as being structured by environmental filtering, with low estimated *m* aligning with expectations of low dispersal. However, the low estimates of *v* and *s_E_* are somewhat surprising, given their documented pattern of single-island endemism (Parent & Crespi 2006). In this case, unmodeled habitat heterogeneity, which is known to be an important predictor of snail diversity (Parent & Crespi 2006), could artificially deflate estimates of *v* and *s_E_* by integrating over unaccounted for local heterogeneity in species trait-environment relationships. Finally, because the Australian tree communities are plot-level samples from smaller scales representing semi-isolated habitat patches and not true insular systems we expect their parameter estimates to deviate from those of true island assemblages. This is in agreement with the finding that these tree communities are all far from equilibrium (Rossetto *et al.* 2015). Specifically, our approach estimates that the system is characterized by moderate *m,* and high *v* and *s_E_* estimates which indicate that local turnover, in the context of a selective environment, is important and ongoing. Additionally, considering the fit of the tree data to a smaller α, the sample abundance in the scaled model and the (unobserved) 'true' abundance that better reflects the effective population size are more similar for trees than for the other datasets.

### Future perspectives

As a first approximation of the feedbacks between processes operating at different timescales MESS makes several simplifying assumptions which can be treated as targets for future model improvement. Non-neutral dynamics could incorporate multivariate trait evolution, allow for filtering and competition processes within the same model, and/or allow for mutualistic rather than simply competitive interactions. Modelling more realistic metacommunity processes and patterns, and including more sophisticated measures of diversity such as temporal correlations and environmental matching would allow for expanding beyond the simple local/regional dichotomy. One caveat is that deviations from panmictic population structure will distort model selection and parameter estimation during MESS inference. For example, cryptic population structure will reduce *S* and inflate metrics of genetic diversity, which could bias MESS inference to prefer non-neutral models, within which these features are common hallmarks. Another special consideration is the variance in the rate at which Λ changes with respect to time as measured in generations. Specifically, the neutral approach to equilibrium is much slower (with respect to numbers of generations) than either of the non-neutral models, potentially confounding comparisons between models at fixed values of Λ. This also highlights the need for a more robust measure of equilibrium, which can account for processes across timescales.

Another approximation is the use of the rescaled Wright-Fisher coalescent process to generate the community-wide population genetic predictions of the forward-time Moran birth/death process. Yet future advances could make use of the powerful new tree-sequence recording (Kelleher *et al.* 2018; Haller & Messer 2019) to more accurately and flexibly match the full demographic and abundance history of each species with its respective underlying population genetic history. Although here we modeled a single locus per species to match the barcode and metabarcode data that are emerging from high-throughput ecological sampling efforts, implementing tree-sequence recording methods could also allow for flexible downstream options to incorporate spatial information associated with genetic geo-reference databases (Lawrence *et al.* 2019).

### Conclusion

The MESS model unifies the study of biodiversity by linking ecological and evolutionary theory across three disparate timescales within an individual-based, mechanistic framework. The model generates explicit temporal predictions of community-scale data across these three diversity axes (species richness and abundance, population genetic diversity, and trait variation), spanning equilibrium and non-equilibrium conditions, and allowing for stochasticity along a continuum of scenarios ranging from pure ecological neutrality, to strong ecological interactions and/or environmental filtering. To complement the MESS model simulations, our implementation includes an extensive suite of ML tools for performing model selection and parameter estimation from observed data, and plotting routines for visualizing and evaluating results. This unified mechanistic model provides a general framework for hypothesis testing and biodiversity data synthesis, enabling the generation of multi-dimensional forecasts and test parameterized hypotheses about the historical and future processes driving biodiversity patterns from small-scale intensively sampled plots, to islands *sensu lato*, to regional and sub-continental scales. With our approach we were able to identify whether real communities were near equilibrium or not, and the eco-evolutionary processes underlying those dynamics. For example, despite the near-equilibrium state of both spider and beetle communities on islands, we discovered that the approach to these equilibria were different, with spider communities assembling largely by immigration, compared to the more prominent role of speciation in weevil communities. This confirms suspected, but as of yet untested, hypotheses from other island arthropod systems (Rominger *et al.* 2016) that can only now be rigorously evaluated. We were also able to pinpoint the mechanistic causes (turnover and environmental filtering) of non-equilibrium in the tree communities. Finally, our analysis of Galapagos snails highlight areas for future improvement in modeling more fine scale environmental heterogeneity and its impact on filtering and speciation.

## Acknowledgements

This paper is a product of the working group sEcoEvo - Biodiversity Dynamics: The Nexus Between Space & Time, which was kindly supported by sDiv, the Synthesis Centre of the German Centre for Integrative Biodiversity Research (iDiv) Halle-Jena-Leipzig and the Santa Fe Institute supported additional working group meetings. We thank John Chase, Catherine Graham, Jacopo Grilli, Joaquín Hortal, Petr Keil, Tiffany Knight, Angela McGaughran, and Brian McGill for useful conversations. Funding was provided by grants from FAPESP (BIOTA, 2013/50297-0 to MJH and AC Carnaval), NASA through the Dimensions of Biodiversity Program (DOB 1343578) and the National Science Foundation (DEB-1253710 to MJH; DEB 1745562 to AC Carnaval; DBI 1927319 to AJR). IO was supported by the Mina Rees Dissertation Fellowship in the Sciences provided by the Graduate Center of the City University of New York. MR was supported by the Bioinformatics and Computational Biology Fellowship through the Institute for Bioinformatics and Evolutionary Studies at the University of Idaho. AJR was supported by the Santa Fe Institute Omidyar Fellowship. JR was supported by fellowships from the Natural Environment Research Council (NERC) (NE/I021179, NE/L011611/1). RSE was supported by an NWO-VICI grant. This work is a contribution to Imperial College’s Grand Challenges in Ecosystems and the Environment initiative, through JR.

